# Detection and evaluation of copy number variation using both linked-read and short-read sequencing in New Zealand dairy cattle

**DOI:** 10.64898/2026.04.20.718595

**Authors:** Yu Wang, Tony Nugroho, Thomas JJ. Johnson, Christine Couldrey, Bevin L. Harris

## Abstract

In recent years, genetic studies have made significant progress in identifying single-nucleotide polymorphisms (SNPs) associated with cattle health and production traits. However, it is still challenging to identify and validate more complicated forms of variation, such as copy number variation (CNV) and other types of structural variation (SV). In this study, SV regions were identified using 37 New Zealand dairy cattle with linked-read sequence data. A transmission-based framework was used to validate these variants at the population scale. 62,438 putative autosomal SV regions were identified with the LongRanger pipeline following the 10x Genomics recommendations. Copy number states for these regions were subsequently estimated via a read-depth based genotyping method using CNVpytor in a population-representative cohort of 2306 animals using Illumina short-read sequencing technology. Mendelian inheritance of copy number states was assessed using linear mixed models incorporating pedigree information, and transmission levels were used to quantify the biological validity of each CNV region. Transmission levels ranged widely, with a mean of 0.5162 across all regions, where higher transmission levels were proportionally enriched for larger SVs. A total of 7218 CNV regions exhibited high transmission levels (>0.9), indicating strong evidence of inheritance. Among these, 7136 overlapped CNV regions reported in one or more public datasets, while 82 high-confidence regions represent previously unreported variants. High-transmission CNV regions tended to show clear, discrete inheritance patterns in trio families, providing the biological evidence that these CNVs are inherited within the population. Together, these results demonstrate that integrating linked-read sequencing with population-scale transmission-based validation provides a robust framework for identifying high-confidence CNV regions. This catalogue of validated CNV regions represents an important resource for downstream functional analyses and the incorporation of structural variation into genomic selection and breeding programs.

## Introduction

Structural variation (SV) represents a major source of genetic diversity and contributes substantially to phenotypic variation in mammals, complementing the effects of single-nucleotide polymorphisms (SNPs) in shaping complex traits (Zhang et al., 2009; Chen et al., 2024). Among different classes of SVs, copy number variations (CNVs), defined as genomic gains and losses typically exceeding 50 bp, are of particular interest due to their direct influence on gene dosage (Rice and McLysaght, 2017) and regulatory elements (Haraksingh and Snyder, 2013). CNV regions have been shown to affect gene expression and biological pathways, which relate to numerous economically important traits (Liu, 2025). In dairy cattle, CNV regions have been associated with production, fertility, health, and longevity traits, highlighting their potential importance for both biological discovery and applied breeding programs (Butty et al., 2021; Oliveira et al., 2024; Ladeira et al., 2025).

Despite their functional relevance, CNV regions remain challenging to detect and interpret accurately. Most large-scale cattle breeding programs rely on SNP-based genotyping platforms, where analytical pipelines and genomic prediction models are well established (VanRaden, 2020). In contrast, accurate CNV detection typically requires whole-genome sequencing and computationally intensive analyses, with performance strongly influenced by accuracy of genome mapping and sequencing depth (Liu et al., 2024). Read-depth–based approaches are scalable but prone to false positives and imprecise copy number estimates, particularly in GC-biased or highly repetitive genomic regions (Bickhart et al., 2012; Duan et al., 2013). As a result, there is no consensus on the best method of CNV identification and many detected CNV regions lack robust biological validation, which limits their utility in downstream analyses and applications in the breeding programs. In cattle, studies comparing sequencing platforms have demonstrated that long-read technologies identify substantially more SVs than short-read approaches, particularly for larger variants (Gao et al., 2022). Long-read sequencing technologies provide an improved resolution for CNV discovery by capturing long-range genomic information and enabling more accurate breakpoint inference (Lin et al., 2023). However, the cost, data volume, and limited throughput of these technologies currently preclude their routine deployment at the population level (Nguyen et al., 2023).

An additional challenge in CNV analysis arises from population structure, breed difference, and reference genome bias. Structural variation is increasingly recognised as partly breed-specific, reflecting demographic history, selection pressures, and divergence between sampled individuals and the reference genome (Chen et al., 2021). Studies have reported significant differences in detected number of CNV regions, size distribution, and genomic location between major dairy breeds such as Holstein-Friesian and Jersey, affecting both discovery power and cross-study concordance (Reynolds et al., 2018). Reference genomes derived from a single breed may further bias detection toward variants present in the reference haplotypes, reducing sensitivity for structurally divergent regions in other populations (Butty et al., 2020). These factors complicate the interpretation of CNV catalogues across different or admixed dairy cattle populations.

Given these constraints, robust validation strategies are essential to distinguish biologically meaningful CNV regions from technical artefacts. Pedigree-based analyses exploiting Mendelian transmission provide a powerful framework for evaluating the biological validity of CNV regions at a population level. Couldrey et al., (2017) demonstrated that CNV regions showing consistent inheritance patterns within families are more likely to represent true structural variants than those detected solely based on read-depth or positional overlap. Transmission-based validation therefore offers a biologically grounded approach that complements discovery-oriented methods and helps bridge the gap between structural variant research and practical breeding applications.

In this study, we integrated the linked-read sequencing–based SV discovery and copy number estimation using short-read sequencing data with population-scale validation in New Zealand dairy cattle. By combining high-resolution discovery in a limited number of animals with transmission-based evaluation across a large, representative cohort, we aim to generate a high-confidence catalogue of CNV regions with strong evidence of inheritance.

## Material and methods

### Linked-read sequencing and structural variants discovery

Linked-read sequencing of 37 New Zealand dairy bulls (18 Holstein-Friesian, 17 Jersey, and 2 Holstein-Friesian Jersey crossbred) was undertaken following the standard protocols by 10x Genomics (Marks et al., 2019). The details of the sequencing procedure were described in Keehan and Couldrey (2017). The long-range context is provided by using microfluidics to segment and barcode high-molecular-weight DNA without needing true long reads. We aligned the reads using the LongRanger (v2.2.2) pipeline following 10x Genomics recommendations. All reads were mapped against the cattle genome reference ARS-UCD 1.2 (Rosen et al., 2020) and underwent subsequent SV detection using the analyze_sv_calls function from the LongRanger pipeline with the recommended settings. SVs were considered for further analysis if they were larger than 50 bp in length, consistent with common definitions of structural variation. Quality control measures excluded any detected regions that lacked start or end positions or were located outside autosomes. After the data cleaning, the called SV regions were then merged across individuals based on genomic coordinate overlap using the bcftools – merge command (Li, 2011), generating a set of non-redundant SV regions.

### Short-reads sequencing and CNV genotyping

Read-depth-based CNV genotyping analysis was undertaken in a population of 2306 animals representing the population structure of New Zealand dairy cattle, which included 730 Holstein-Friesian, 468 Jersey, 1069 Holstein-Friesian × Jersey crossbred, 3 Ayrshire animals, and 36 animals from mixed or minor breeds. They were sequenced on Illumina HiSeq 2000 instruments targeting 100bp paired-end reads. The raw sequence data were mapped to the ARS-UCD1.2 bovine reference genome (Rosen et al., 2020) using the Burrows-Wheeler alignment algorithm version 0.7.17 (Li, 2013). The copy number states of the identified regions were then estimated in each of the short-read sequenced animals using the genotyping function from CNVpytor version 1.3.1 (Suvakov et al., 2021), a Python version of its previous software, CNVnator (Abyzov et al., 2011). Standard CNVpytor quality control and GC bias correction were applied prior to copy number estimation. The read-depth signals were extracted and compared to the expected distribution for different copy number states for each provided region by dividing the genome into segments of 200bp. For each putative SV region, copy number estimates from constituent bins were aggregated to assign a single copy number state per animal per region. Due to the lack of SV type annotation in the LongRanger pipeline, we implemented a classification method based on the copy number status. The state of each region was categorized into loss, normal, and gain state via copy number estimates per individual. Because copy number estimates derived from read depth are continuous and subject to technical variation, thresholds allowing for uncertainty around the diploid state were applied. Copy number values lower than or equal to 1.5 were classified as losses, values greater than or equal to 2.5 as gains, and intermediate values as normal (Friedrich et al., 2020). The frequency of each state was calculated across all individuals for all regions. Regions were classified as deletion or duplication CNV regions when the corresponding state was present in at least 50% of individuals. Regions showing both loss and gain in at least 5% of individuals were classified as complex CNV regions. Regions with low-frequency copy number changes (<5%) were classified as rare CNV regions, while regions without a dominant state and exhibiting high variance were labelled as uncertain.

### Transmission-based validation

Mendelian inheritance of these copy numbers was then estimated using a univariate animal model, which was described in detail in Couldrey et al., (2017). The model is specified as:

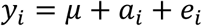

where 𝑦_𝑖_ is the estimated copy number state of a given region for the individual 𝑖, 𝜇 is the overall mean, 𝑎_𝑖_ is the additive genetic effect of the individual 𝑖 and 𝑒_𝑖_ is the residual error. The additive genetic effects are assumed to follow: 𝑎 ∼ 𝑁 (0, 𝐴𝜎_a_^2^), where 𝐴 is the relationship matrix derived from pedigree traced for 3 generations and 𝜎_α_^2^is the additive genetic variance. The residual errors are assumed to be independently and normally distributed: 𝑒 ∼ 𝑁 (0, 𝐼𝜎_𝑒_^2^), where 𝐼 is the identity matrix and 𝜎_𝑒_^2^ is the residual variance. Transmission level therefore reflects the proportion of variance in estimated copy number attributable to additive genetic effects. Transmission level ranged from 0 to 1, with 0 indicating the region was a false positive discovery or a sequence artifact and 1 indicating the copy number is inherited following Mendelian inheritance. The APEX linear model suite (www.ghpc.ai) was used to efficiently solve these equations. Copy number inheritance was additionally examined and visualized in 600 trio families to assess Mendelian transmission patterns.

### Comparison with publicly available CNV datasets

Four sets of previously reported CNV datasets of dairy cattle were considered for comparison with the copy number variants detected in this study. Table S1 provides an overview of these studies, including data type, sample size, breeds, analytical methods, and database availability. CNV regions archived in the Database of Genomic Variants Archive (DGVa) were obtained from eight studies and remapped to the ARS-UCD1.2 assembly to ensure consistency with the reference genome used in this study. Additional CNV datasets were obtained from Lee et al., (2023), Bhati et al., (2023) and Grant et al., (2024). Overlapping CNV regions among datasets were identified using the *GenomicRanges* R package (Lawrence et al., 2013). An overlap of at least 10 bp was required to classify equivalence between our detected region with a publicly available region. Genome feature annotations, including gene and exon coordinates, were downloaded from Ensembl (Bos_taurus.ARS-UCD1.2.110.gtf.gz) for downstream annotation of overlapping regions.

## Results

In this study we utilized linked-read sequencing to discover potential SV regions. These putative SVs regions were then validated in short-reads sequenced animals by evaluating the copy number status and the estimating the transmission levels. Together, these analyses define a genome-wide set of CNV regions with strong population-level evidence of inheritance.

### Structural variants inference

All 37 linked-read sequenced animals have an average sequence depth between 29x and 43x, except for a bull, “Esteem,” which has a sequence depth of 68.89x with the highest number of detected reads. The relationship between average sequence depth and number of structural variants detected can be seen in Supplemental Material Figure S1. While some variation is observed, SV discovery does not scale linearly with the sequence depth, and three animals with extreme values are highlighted. The number of reads detected among animals ranged between 634,723,606 and 1,573,807,188 (Esteem) and the mapping rate ranged between 83.75% and 96.95%. On average, 850,231,229 reads were generated by 10x genomics, representing 39.7x coverage of the genome. Approximately 7000 SVs were discovered in each animal. One bull (“Lonestar”) showed substantially fewer SVs detected (5,916) and lower mapping rate (83.75%) than other animals, resulting in fewer shared SV regions (∼2,000), consistent with reduced discovery power. On average, the number of regions intersecting among individuals was around 4000. Three animals that have over 5000 overlapping regions originated from a duo family. The number of overlapping SV regions between pairs of individuals sampled were also investigated within the same breed and between different breeds among all 37 animals, which can be seen in Figure 1. The distributions show consistently higher overlap among individuals from the same breed than between breeds. The number of overlapping regions between Lonestar and others was around 2000 across all breed combinations, which is significantly lower than the other comparands.

**Figure 1.**
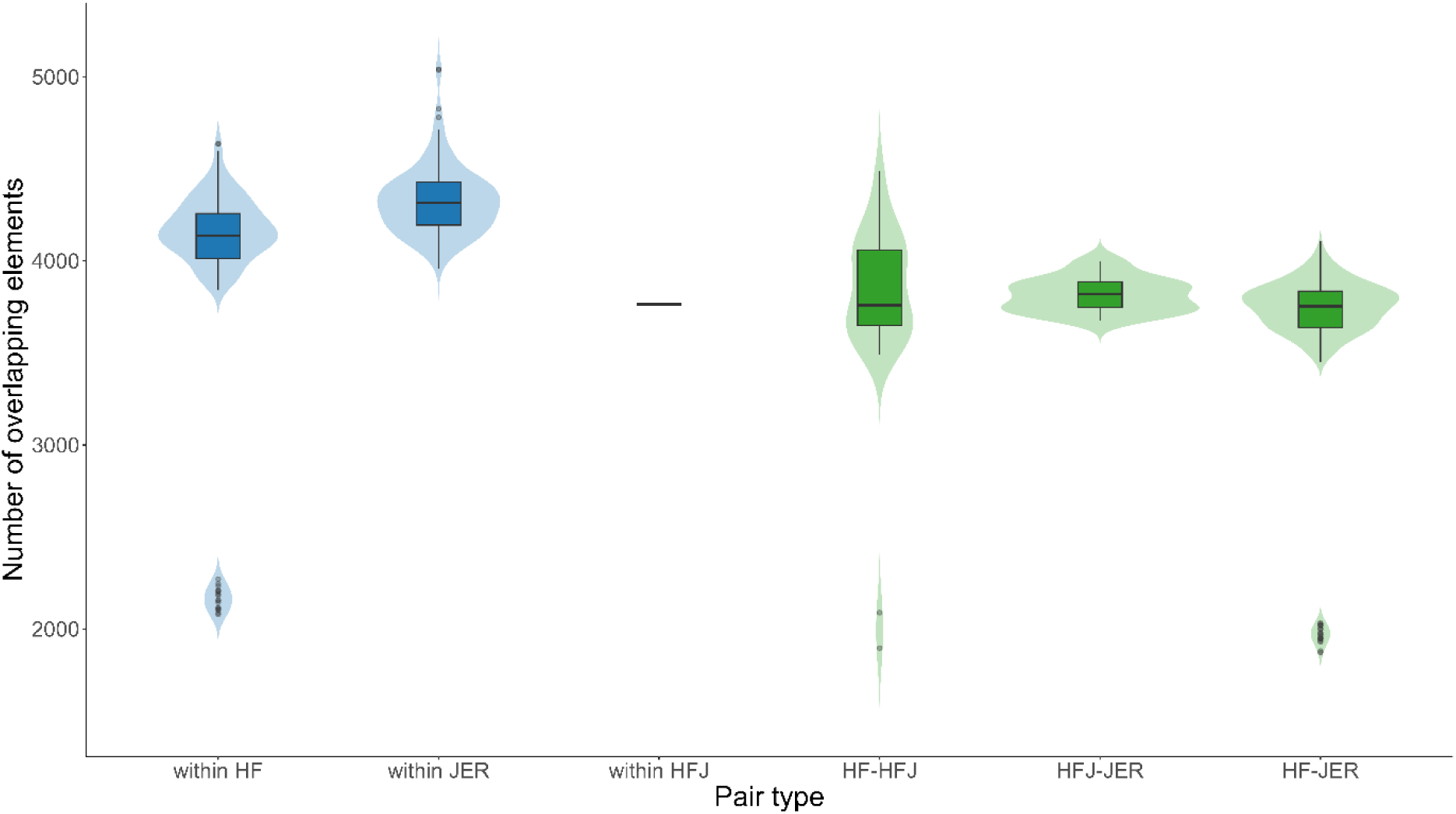
Distribution of overlapping SV regions among pairs of individuals within and between breed groups.^1^ ^1^Blue violins represent within-breed comparisons, while green violins represent between-breed comparisons. ^2^The breed groups are: HF: Holstein-Friesian; JER: Jersey and HFJ: Holstein-Friesian Jersey cross.

After merging the output from the LongRanger pipeline for all animals, 62,438 putative autosomal SVs over 50bp were identified from 37 linked-read sequenced animals. The relationship between autosomal chromosome length and the number of detected SV regions is shown in Figure S2. Each point represents one chromosome, and a strong positive correlation (r=0.843) was observed between chromosome length and the number of detected CNV regions. Chromosome 1, the longest chromosome of the cattle genome, has the most SVs detected with 3498 SVs. Meanwhile, chromosome 25, the shortest chromosome of the cattle genome, has the least SVs detected with 903 SVs. Overall, the cumulative length of SV regions is 33,002,890 bp which accounts for 1.33% of the total length of the genome. Figure 2 presents the size and CNV type distribution of putative SV regions detected across the 37 linked-read sequenced animals. The size of the detected SVs ranged between 50bp and 29,904bp, with an average of 1854bp. Most detected SVs fall into smaller size categories, with progressively fewer SVs detected as size increases.

**Figure 2.**
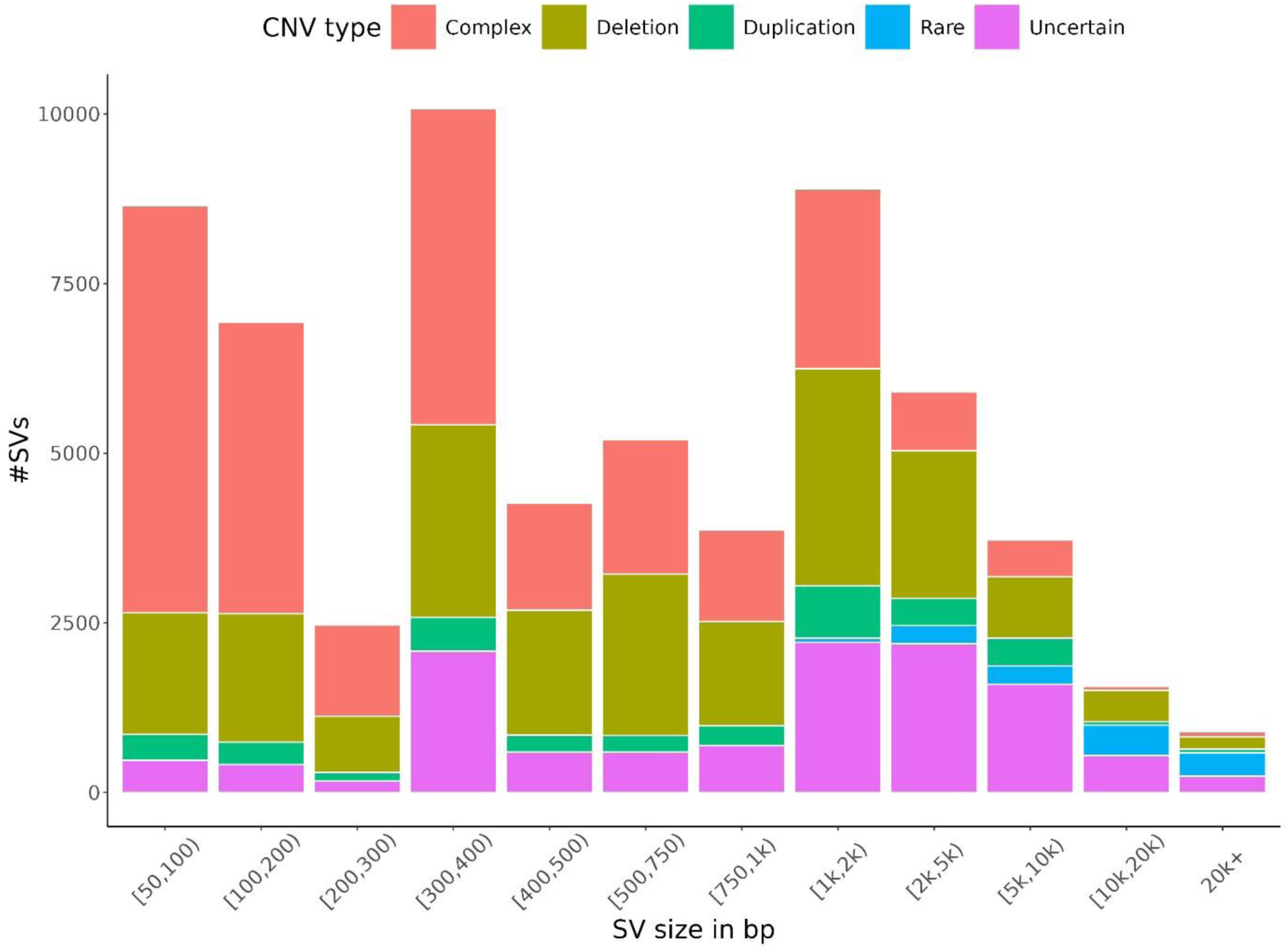
Size distribution of putative structural variant regions detected across 37 linked-read sequenced animals with variant type classification based on the copy number statuses of the short-read sequenced animals.

### Copy number evaluation and transmission level estimation

The CNV type for each detected SV region was determined by estimating the proportion of individuals exhibiting gain, loss, or normal states. In summary, most regions were classified as complex (40.69%), deletion (32.07%), or uncertain (18.88%). Duplication accounted for 6.14% of detected regions, while 2.23% were considered rare. The average sizes of regions across most categories were comparable, except for the rare category, which exhibited significantly larger average sizes at 12,895 bp (see Table 1).

**Table 1.** Number, average size and average transmission level of CNV regions of each SV type.

|  | Total number | Size in bp (mean±sd) | Transmission level (mean±sd) |
| --- | --- | --- | --- |
| Deletion | 20,022 (32.07%) | 1664±3345 | 0.714±0.241 |
| Duplication | 3828 (6.13%) | 2201±4071 | 0.537±0.202 |
| Complex | 25,406 (40.69%) | 740±1912 | 0.398±0.272 |
| Rare | 1391 (2.23%) | 12,895±8346 | 0.241±0.198 |
| Uncertain | 11,791 (18.88%) | 3164±4757 | 0.460±0.311 |

To assess the biological validity of the linked-read–discovered SVs, transmission levels were estimated using population-scale short-read sequencing and pedigree information. The 62,438 putative SV regions discovered in the linked-read sequence data showed a wide range of transmission levels on the 2306 short-read sequenced animals (see Figure 3). The average transmission level was 0.5162 with a standard deviation of 0.3016. CNV regions were fairly evenly spread across each transmission level bin split by 0.1, with the counts in these bins ranging from 4,949 to 7,629. This represents between 7.93% and 12.22% of the total number of CNV regions (refer to Table 2). While small CNV regions are numerically dominant overall, Figure 4 shows that higher transmission levels are proportionally enriched for larger CNV regions, whereas lower transmission level bins are dominated by smaller CNV regions. Notably, regions designated as deletions demonstrated a higher transmission level relative to other categories, averaging 0.71 (see Table 1).

**Figure 3.**
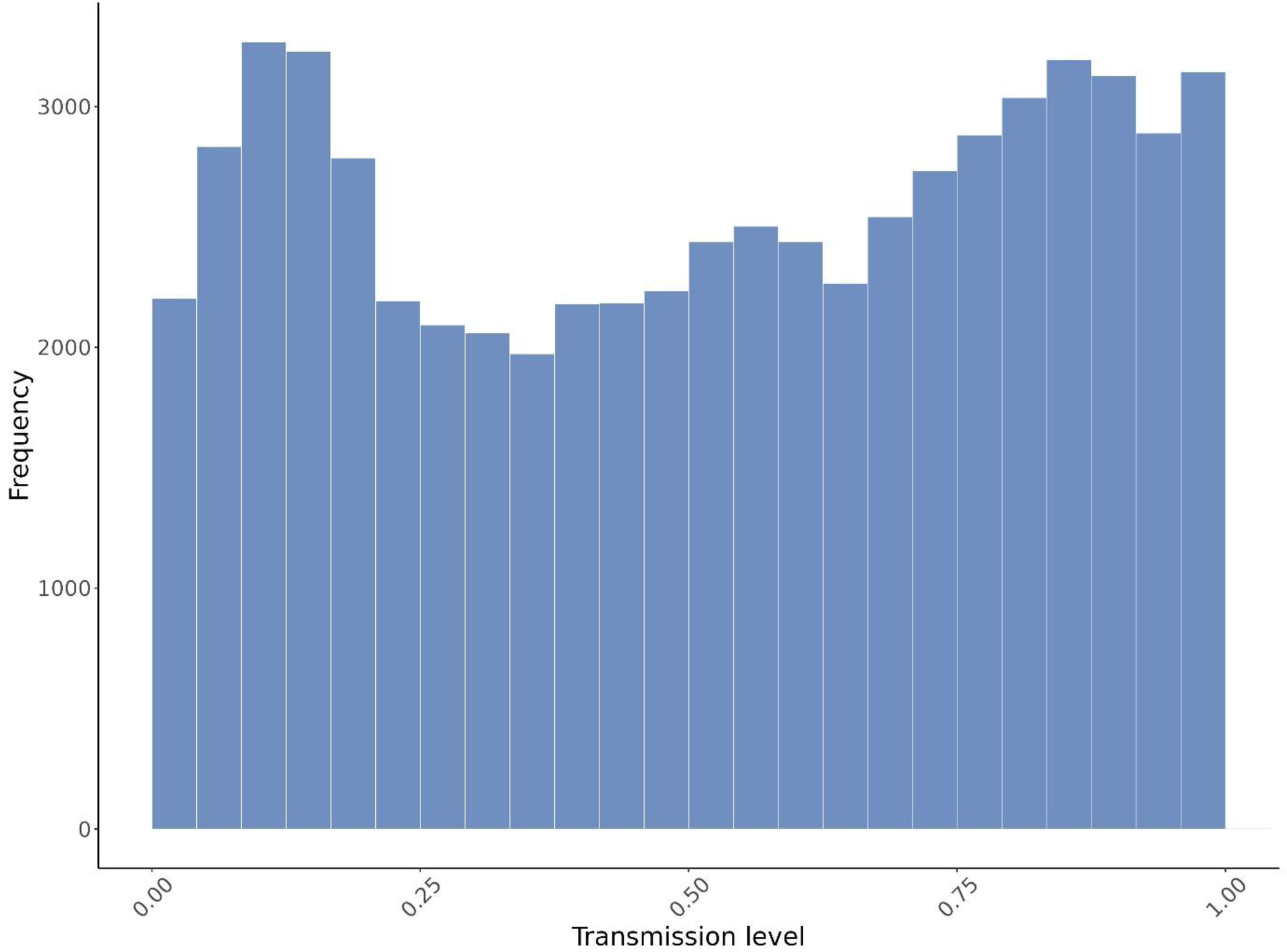
Distribution of transmission levels for putative structural variant regions detected by linked-read sequencing and evaluated in 2,306 short-read sequenced dairy cattle.

**Figure 4.**
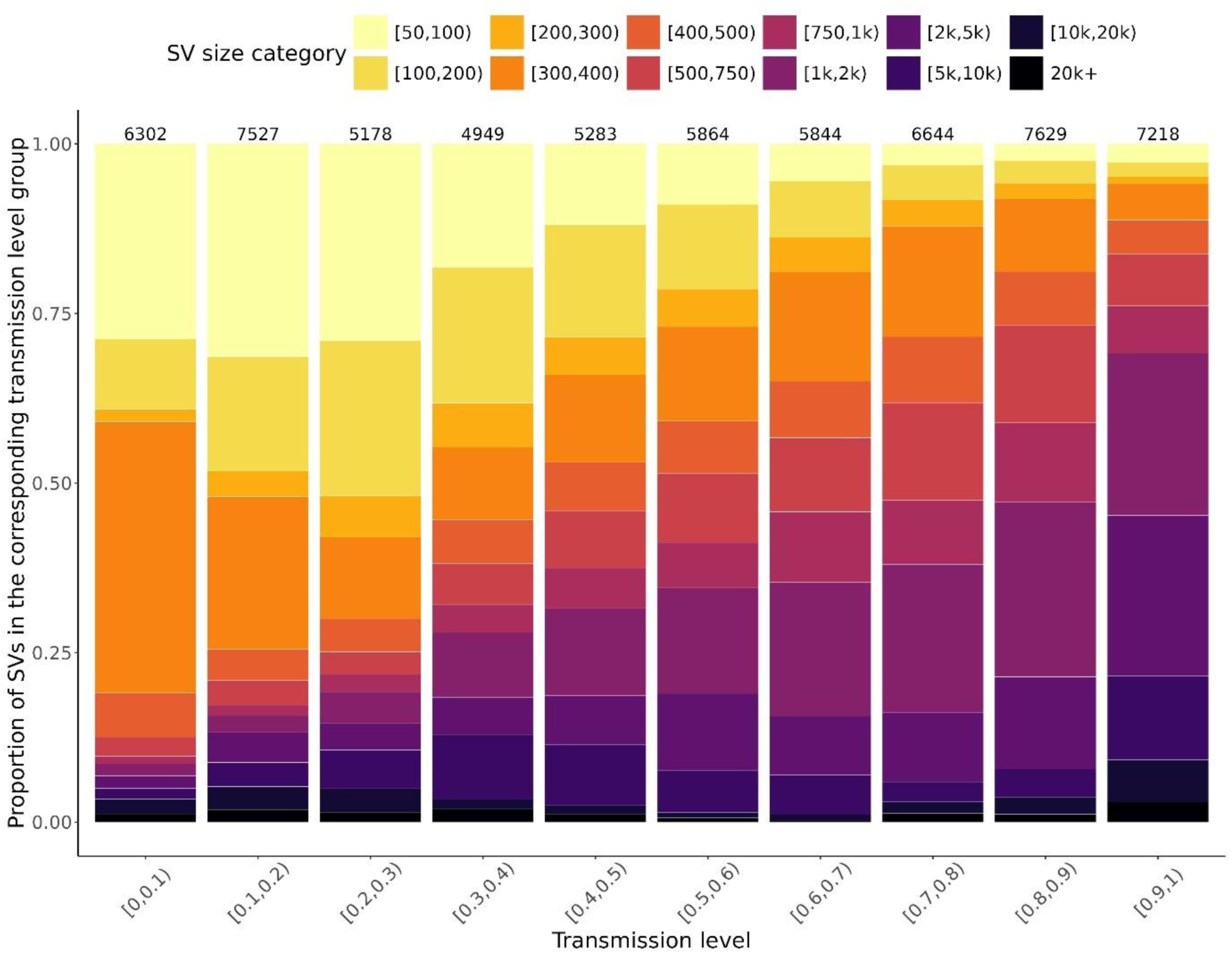
Proportional distribution of SV size categories within each transmission-level bin.^*^ ^*^The number of SV in each transmission level category is above each bar. Each stacked bar sums to 1 and represents the relative contribution of each SV size category conditional on transmission level.

**Table 2.** Number and proportions of CNV regions in each transmission level group.

| Transmission level group | Number of CNV regions | Proportion of CNV regions |
| --- | --- | --- |
| [0,0.1) | 6302 | 10.09% |
| [0.1,0.2) | 7527 | 12.06% |
| [0.2,0.3) | 5178 | 8.29% |
| [0.3,0.4) | 4949 | 7.93% |
| [0.4,0.5) | 5283 | 8.46% |
| [0.5,0.6) | 5864 | 9.39% |
| [0.6,0.7) | 5844 | 9.36% |
| [0.7,0.8) | 6644 | 10.64% |
| [0.8,0.9) | 7629 | 12.22% |
| [0.9,1) | 7218 | 11.56% |

CNV regions with higher transmission levels tended to follow a multimodal distribution of copy number estimates among animals and the inheritance can be clearly observed in the trio families. In Figure 5, we illustrated a CNV region with a transmission level of one. This CNV was selected as a representative example of high-transmission deletions observed genome-wide. From the distribution of the copy number of this CNV region (Figure 5a), most data points clustered around integer CNV values (0, 1, 2). CNV values of 0 and 1 are more frequent than 2, indicating this copy number variant is likely to be a deletion. Figure 5b illustrates copy number inheritance across sire, dam, and offspring for the same CNV region across 600 trio families. The clustering at integer values suggests that copy number variations are mostly discrete among all individuals. A positive correlation can be observed between parents’ copy numbers and their offspring’s copy number. When both sire and dam have a copy number of 0, their offspring also tend to have a copy number of 0, indicating a zero-copy deletion being inherited by the offspring.

**Figure 5.**
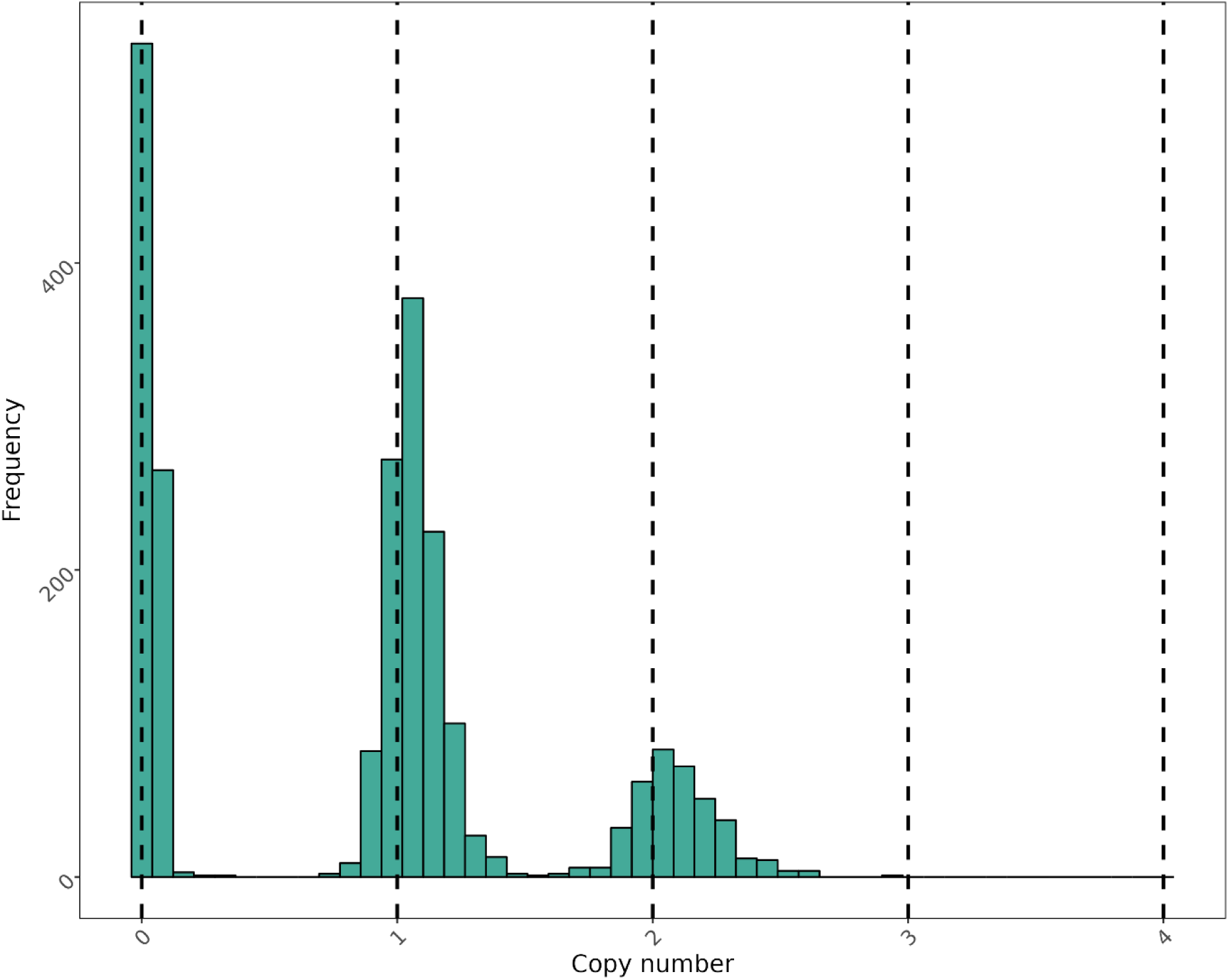
a. Distribution of copy number states for a representative CNV with complete Mendelian transmission (transmission level = 1).

**Figure 5.**
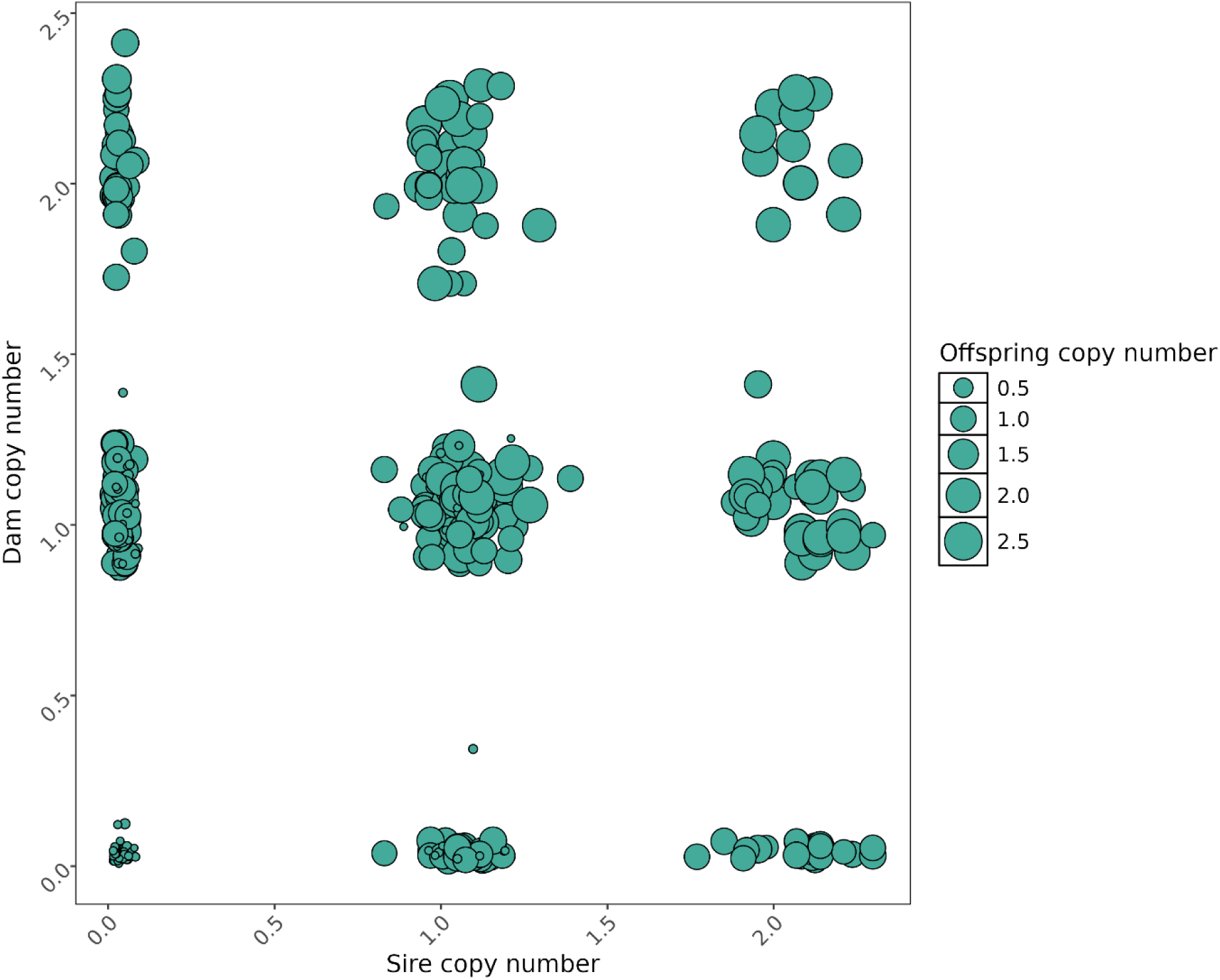
b. Copy number inheritance patterns among sire, dam, and offspring across 600 trio families for the CNV shown in Figure 5a.

### Overlap with other publicly available databases

We compared our putative CNV regions with those in four public datasets. The number of CNV regions reported in each of these public datasets ranged from 13,732 to 65,550 which can be seen in Table 3. A substantial proportion of previously reported CNV regions were rediscovered in our dataset, with over half of regions from DGVa, Lee et al. (2023), and Bhati et al., (2023) overlapping with regions identified in this study. In contrast, overlap with CNV regions reported by Grant et al. (2024) was lower, with only 44.15% of regions using a Menta pipeline and 25.79% of regions using a Smoove pipeline. Inversely, the public dataset that contained the highest number of regions found in this study was DGVa with more than 86% of our regions overlapping. The public dataset that contained the lowest number of regions found in this study was Bhati et al., (2023) with less than 39% of our regions overlapping.

**Table 3.**
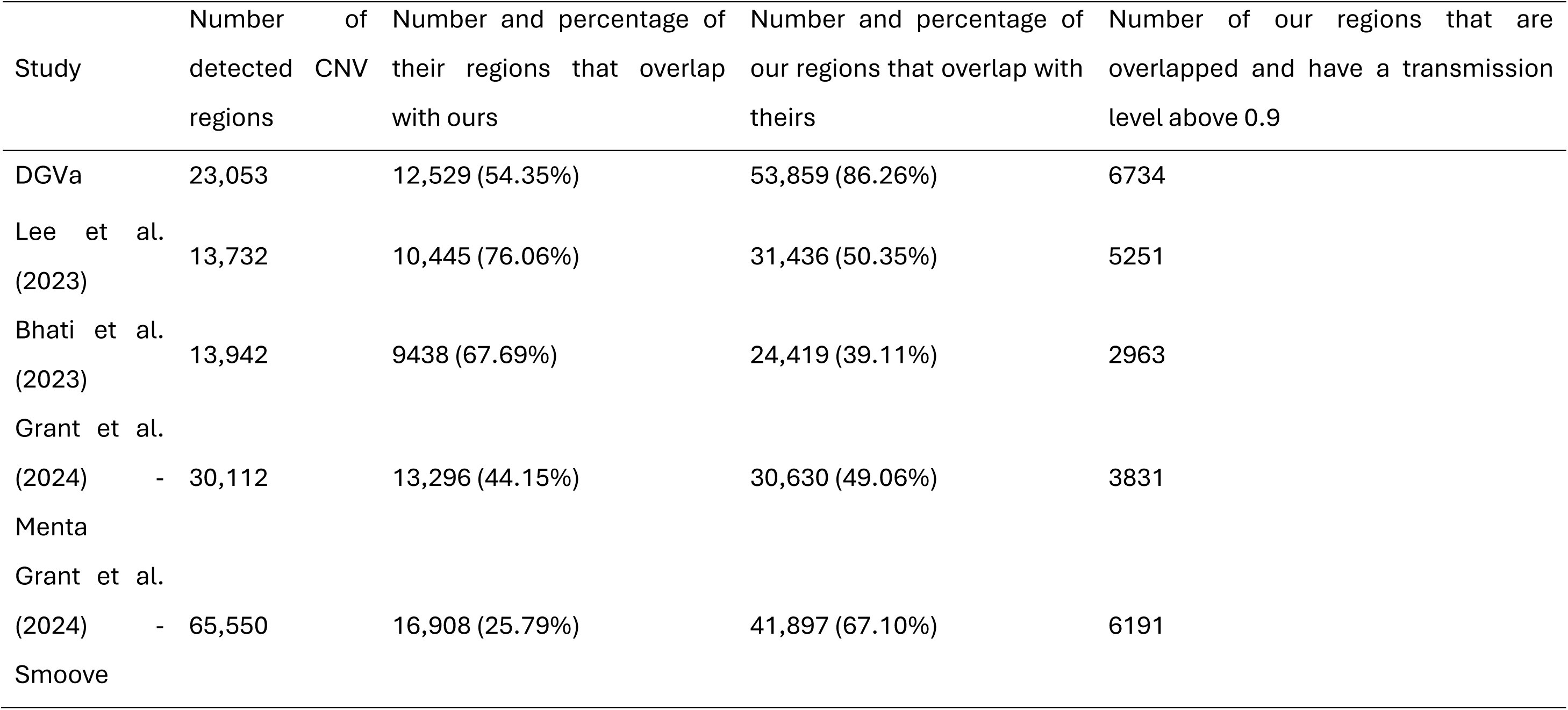
Comparison of CNV regions identified in this study with publicly available CNV datasets of dairy cattle, showing overlap counts and transmission-based validation.

7218 CNV regions found in this study exhibited high transmission levels (> 0.9). Among these, we identified 5267 deletions, 196 duplications, 994 uncertain, 728 complex and 33 rare variants. 82 of these regions represent novel variants that were not reported before, while the remaining regions overlapped one or more published datasets. CNV regions identified consistently across multiple studies tended to show higher transmission levels. 6734 high-transmission CNV regions overlapped regions in DGVa, representing the largest concordant set among all comparisons, whereas fewer high-transmission CNV regions were overlapped with Bhati et al., (2023). 1561 of the regions that have a high transmission level are locating on the region of genes, in which 357 of them were located on an exon.

## Discussion

In this study we developed a pipeline combining SV discovery using linked-read sequence data, copy number estimation and population scale transmission-based validation using short-read sequencing data in dairy cattle. Firstly, the SV regions were identified in 37 animals with linked-read sequence data. The copy number status for these regions was then assessed on 2306 animals with Illumina short-read sequencing information. We then evaluated the transmission level for each region to assess biological evidence for validation. These regions were also compared with several publicly available databases.

Structural variant discovery has commonly relied on whole-genome sequence data, yet the effectiveness of the analysis depends on the chosen sequencing technology with each offering its own set of trade-offs (Lavrichenko et al., 2021). Conventional short-read whole-genome sequencing provides high base accuracy and scalability (Zhao et al., 2021), but its limited read length constrains the ability to resolve larger SVs, particularly those spanning repetitive or hard-to-map regions. In addition, SV detection from short-read data relies largely on indirect signals such as read depth, paired-end discordance, and split reads, which are sensitive to mapping artefacts and often yield imprecise breakpoints. These limitations contribute to both reduced sensitivity for smaller variants and an elevated false-positive rate, especially in complex genomic regions. In contrast, long-read sequencing platforms, such as PacBio (Byrska-Bishop et al., 2022) and Oxford Nanopore (Wang et al., 2021), represent the current gold standard for SV discovery. They offer the ability to span entire variants and resolve breakpoints at base-pair resolution. Long-reads consistently outperform short-read approaches in detecting large, complex, and nested SVs, including insertions and rearrangements in highly repetitive regions. However, long-read sequencing remains constrained by higher costs, lower throughput, and practical challenges in generating population-scale datasets, limiting its routine application. In this study, we utilized the linked-read sequencing technology, which partially overcomes the challenges of providing long-range information to short reads and can detect large SVs at a lower cost than long-reads sequencing technology (Spies et al., 2017; Karaoğlanoğlu et al., 2020). While LongRanger itself is no longer actively developed, the transmission-based validation framework presented here is independent of the discovery technology and is directly applicable to SVs detected using long-read, pangenome, or graph-based approaches.

A key challenge in structural variant discovery is distinguishing true inherited variants from technical artefacts generated by sequencing and alignment errors. This challenge is amplified as sequencing technologies with higher discovery sensitivity, such as linked-read sequencing, increase the number of candidate SV regions. In this study, the number of putative SVs identified per animal using linked-read data ranged between 5916 and 7981 among all 37 individuals. These results are comparable to Gao et al. (2022), where 8,315 SVs were detected in an individual with a sequencing depth of 55x. However, the number of detected SV regions are significantly larger compared to the other studies using short-read sequencing technology (see Table 1). These counts substantially exceed those typically reported from short-read only pipelines, which generally detect fewer and smaller SVs due to limitations in breakpoint resolution and mapping ambiguity.

To address the inherent ambiguity of LongRanger’s breakpoint-centric output, we implemented a robust copy-number-based classification pipeline that translates abstract breakpoints into biologically interpretable structural variants. By utilizing depth-of-coverage from short-read data, a conservative thresholding strategy (copy number ≤1.5 for losses and copy number ≥2.5 for gains) was applied. In diploid genomes, the expected baseline copy number for autosomal loci is two. However, read depth–based copy number estimates are continuous values subject to sampling variance, GC bias, mapping bias, and normalization artifacts. Therefore, many studies do not use an exact cutoff at 2.0 but instead define buffered intervals around the diploid state to reduce false positives. Furthermore, stratifying variants into deletion/insertion (≥50% prevalence), rare (≤5% prevalence), and complex (≥5% prevalence) categories enables the distinction between stable polymorphisms and multi-allelic “hotspots” of genomic instability. In addition, we included an “uncertain” category for regions with high variance which serves as an essential quality control filter. In summary, this thresholding approach balances biological plausibility with measurement uncertainty, thereby improving robustness in large-scale CNV classification analyses.

Despite increasing sensitivity of SV discovery technologies, there remains no widely adopted population-scale framework to distinguish inherited CNV regions between technical artefacts and biological evidence. Transmission-level analysis provides a novel and biologically grounded approach to address indeterminate validity of detected CNV regions by directly evaluating whether inferred copy number states behave as heritable genetic variants at the population level. Unlike commonly used validation strategies based on variant size, frequency, or overlap with external databases, transmission-level analysis quantifies the proportion of variance in copy number estimates attributable to additive genetic effects. Variants with high transmission levels therefore show consistent inheritance across generations, whereas regions with low transmission are more likely to reflect mapping artefacts or stochastic read-depth noise. Figure 4 illustrates that although linked-read sequencing enables the discovery of a large number of putative SVs, these variants span a wide range of transmission levels, with a mean close to 0.5. Our results demonstrate that a substantial fraction of linked-read discovered SVs show weak or negligible evidence of Mendelian inheritance, highlighting that even technologies with enhanced sensitivity remain prone to artefactual calls. This observation underscores that linked-read sequencing, while superior to short-reads for SV discovery, still requires robust downstream validation to distinguish true inherited variants from technical noise.

High-transmission CNV regions identified in this study show strong reproducibility across independent datasets, reinforcing transmission level as a reliable benchmark of biological evidence. Among the 7218 CNV regions with transmission levels greater than 0.9, the vast majority overlapped CNV regions reported in one or more public resources, with more than 86% overlapping regions archived in the Database of Genomic Variants Archive and substantial concordance with other large cattle CNV studies. In contrast, CNV regions with lower transmission levels showed reduced overlap with external datasets, suggesting that poor reproducibility reflects technical noise rather than population specificity. The strong composition of high-transmission CNV regions in variants consistently observed across studies supports the principle that reproducibility across populations and platforms is a hallmark of true inherited structural variation. Importantly, the identification of a small set of novel high-transmission CNV regions using these robust inheritance patterns indicates that a lack of prior annotation does not preclude biological authenticity. This depth-based validation is particularly reliable for deletions, which LongRanger detects with high sensitivity by leveraging barcode-overlap signals (Marks et al., 2019). This has also been reflected in the transmission level among all five categories in which deletions had the highest average transmission levels.

Structural variant discovery is inherently influenced by population composition and breed background, reflecting both genetic diversity and reference genome bias. In this study, SVs detected using linked-read sequencing showed clear evidence of breed and population effects. This is consistent with previous reports that CNV number, size distribution, and genomic location vary substantially between dairy breeds such as Holstein-Friesian and Jersey (Reynolds et al., 2018). These differences likely arise from distinct demographic histories, selection pressures, and structural divergence relative to the reference genome. In this study, the linked-read sequenced animals are prominent bulls that have been frequently involved in breeding schemes. Therefore, the population, which includes the short-read sequenced animals, is very similar and closely related to the original animals used to detect SV regions. In contrast, CNV regions that are rare, breed-specific, or poorly represented in the reference haplotypes are more difficult to detect and validate consistently. We found that over 50% of the detected regions overlap with DGVa, Lee et al., (2023), and Grant et al., (2024). However, the lower overlap with Bhati et al., (2023) likely reflects breed composition differences, as their study included Fleckvieh animals, which are genetically more distant from the Holstein-Friesian and Jersey populations analysed here. Similarly, Chen et al., (2021) detected more SVs in Holstein animals compared to Jersey animals. These findings highlight that SV catalogues are not universally transferable across populations and emphasise the importance of population-matched discovery panels and breed-aware interpretation when generating and applying structural variant resources in livestock genomics.

A validated set of high-transmission CNV regions provides new opportunities to extend the genetic architecture captured in animal breeding programs beyond SNP variation. CNV regions with strong Mendelian inheritance are more likely to represent stable additive genetic effects, making them suitable candidates for inclusion in genomic prediction models. Incorporating such CNV regions alongside SNPs may improve the proportion of genetic variance explained, particularly for traits influenced by structural variation that is poorly tagged by SNP markers (Chen et al., 2021). High-confidence CNV regions could be modelled as additional loci, grouped into regional genomic features, or used to refine haplotype-based prediction frameworks. Breed-specific CNV catalogues further enable the capture of population-specific genetic effects, reducing reference bias and supporting more accurate estimation of breeding values. Together, these developments highlight the potential of validated CNV regions to improve the performance of genomic selection.

## Conclusion

In this study, we combined linked-read sequencing–based discovery with population-scale validation to generate a high-confidence catalogue of copy number variation in New Zealand dairy cattle. Linked-read sequencing of 37 animals identified 62,438 putative autosomal CNV regions, which were subsequently genotyped in 2306 animals using short-read whole-genome sequencing. By estimating transmission levels using pedigree-based linear mixed models, we quantified Mendelian inheritance and distinguished biologically meaningful CNV regions from technical artefacts. A total of 7218 CNV regions exhibited high transmission levels (>0.9), most of which overlapped previously reported variants, while a small number represented novel regions. These results demonstrate that transmission-based validation provides a robust and complementary framework for CNV discovery and confirms the utility of integrating linked-read and short-read sequencing for structural variation studies in livestock populations. Our results highlight the influence of population composition and reference genome choice on CNV discovery and validation, consistent with growing evidence that structural variation is partly breed-specific. By providing a scalable, biologically grounded validation strategy, this work helps bridge the gap between structural variant discovery and practical application in genomic selection and lays the groundwork for further improving the biological interpretation of structural variation in cattle genomes.

## Data Availability Statement

The list of CNV regions and associated transmission level estimates generated in this study are provided in the Supplementary Materials, which include genomic coordinates, CNV type, and transmission level estimates used in all downstream analyses and figures. The linked-read sequence data used for structural variant discovery are available upon request. The Illumina short-read sequencing data and pedigree information used for population-scale validation are confidential and are not publicly available. Requests for access to these datasets should be directed to the corresponding author and will be considered subject to appropriate data-sharing and confidentiality agreements.

## Ethics statement

Ethical approval was not necessary for this animal study, as all analyses were conducted using previously collected datasets, in line with applicable local regulations and guidelines.

## Author contributions

YW: Methodology, Investigation, Resources, Formal Analysis, Visualization, Writing – original draft, Writing – review and editing. TN: Investigation, Data curation, Formal Analysis, Software. Writing – review and editing, TJJJ: Methodology, Data curation, Software, Writing – review and editing. CC: Conceptualization, Project administration, Methodology, Supervision, Resources, Writing – review and editing, Funding acquisition. BLH: Conceptualization, Methodology, Writing – review and editing, Funding acquisition.

## Funding

This study received financial support from the NZ Ministry of Primary Industries, SFF Futures Program: Resilient Dairy–Innovative breeding for a sustainable dairy future (grant number: PGP06-17006).

## Acknowledgments

We thank Mike Keehan for contributions to early data preparation. This research was made possible using the supercomputing resources of the New Zealand eScience Infrastructure (NeSI) platform (https://www.nesi.org.nz).

## Conflict of Interest Statement

Y.W., T.J.J.J., and B.H. are employees of Livestock Improvement Corporation (LIC; Hamilton, New Zealand), a commercial provider of bovine germplasm. T.N. and C.C. was affiliated with LIC during the research; subsequent changes in affiliation do not affect the integrity or interpretation of the findings presented. The authors declare that the research was conducted in the absence of any commercial or financial relationships that could be construed as a potential conflict of interest.

---

**Figure S1.**
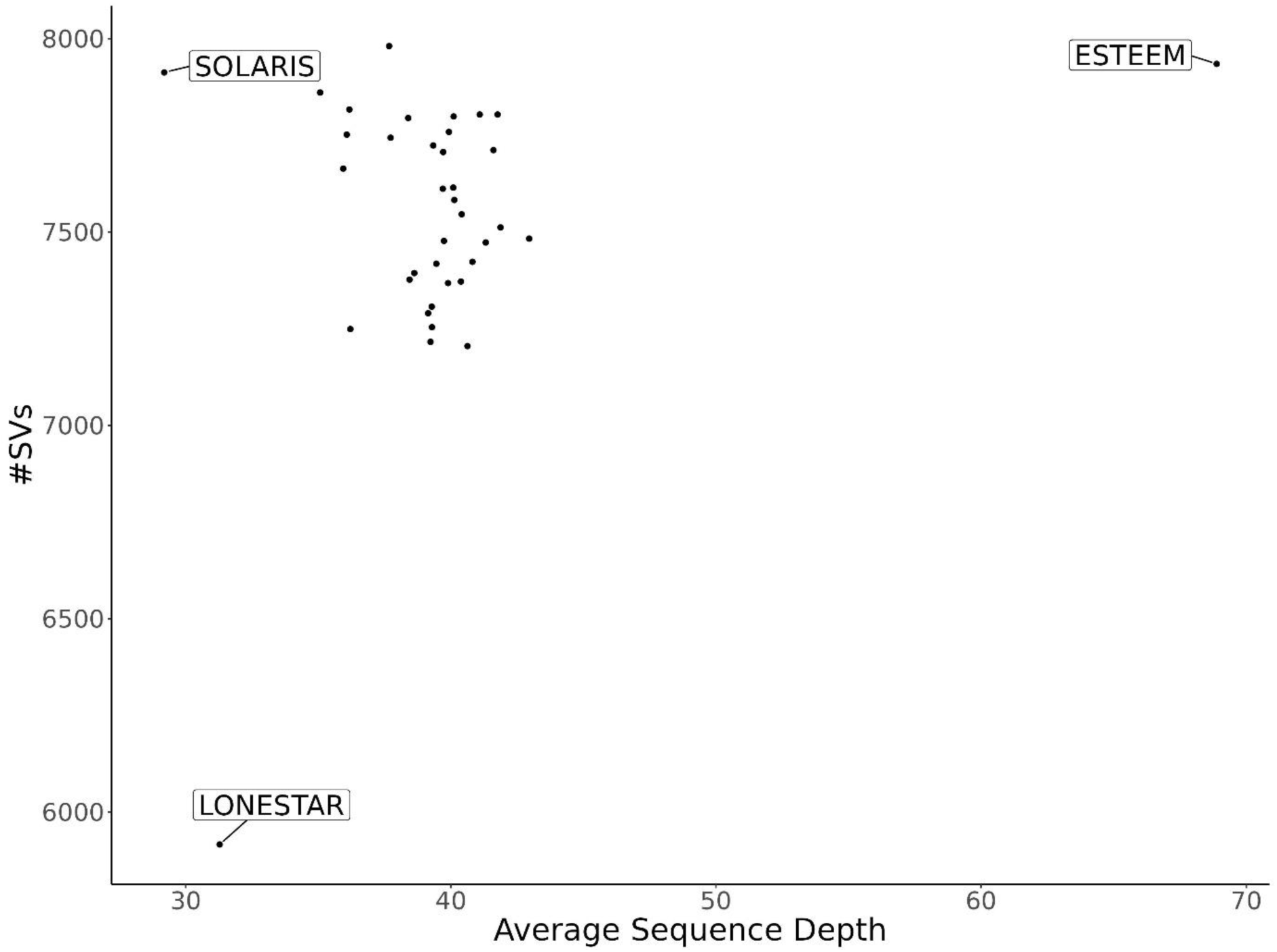
Relationship between the average sequencing depth and the number of detected structural variant regions detected in 37 linked-read sequenced animals.* *Three bulls with highest/lowest sequence depth and number of detected structural variants are highlighted

**Figure S2.**
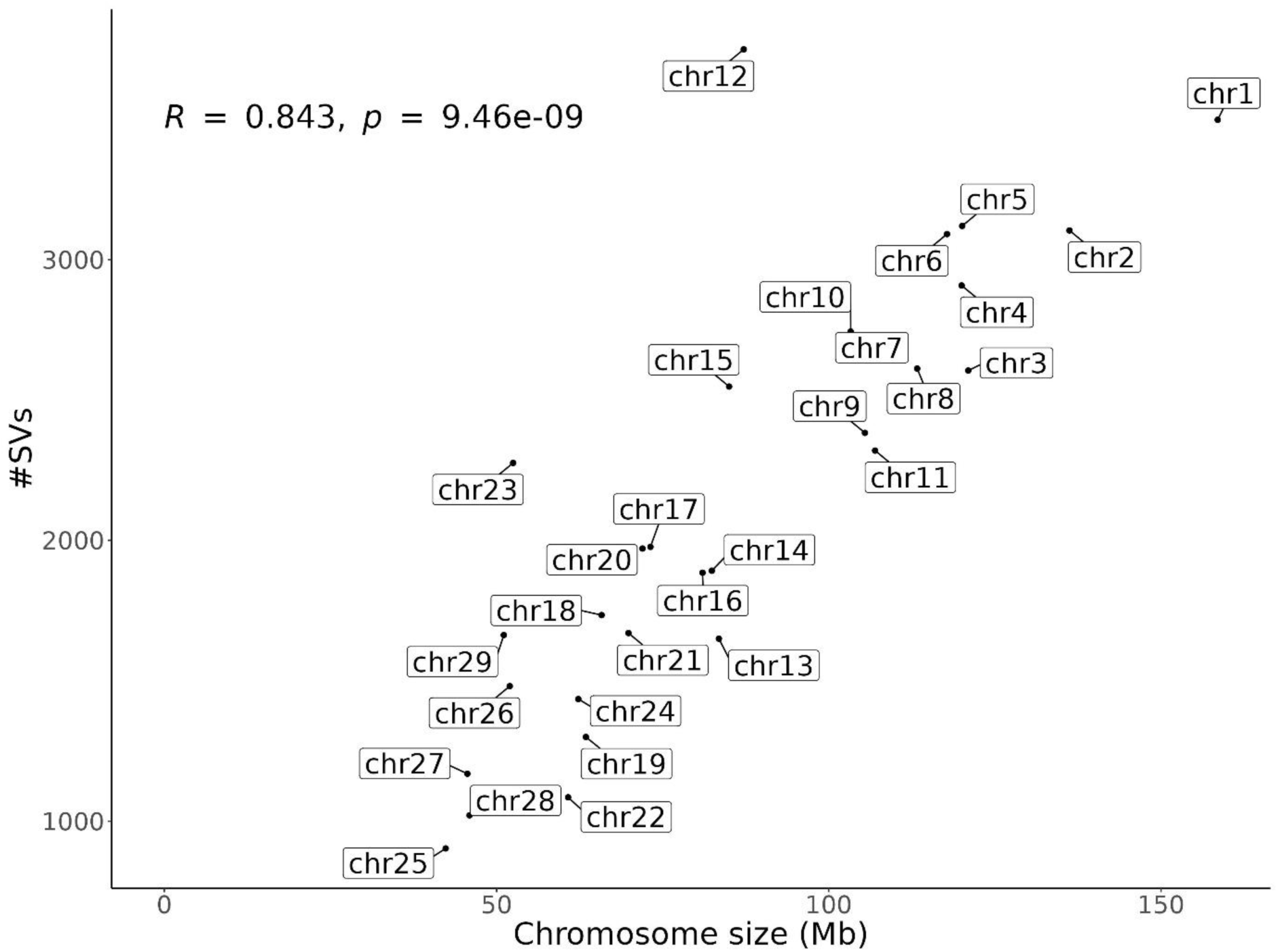
Relationship between chromosome length and the number of detected CNV regions across the cattle genome.

**Table S1.**
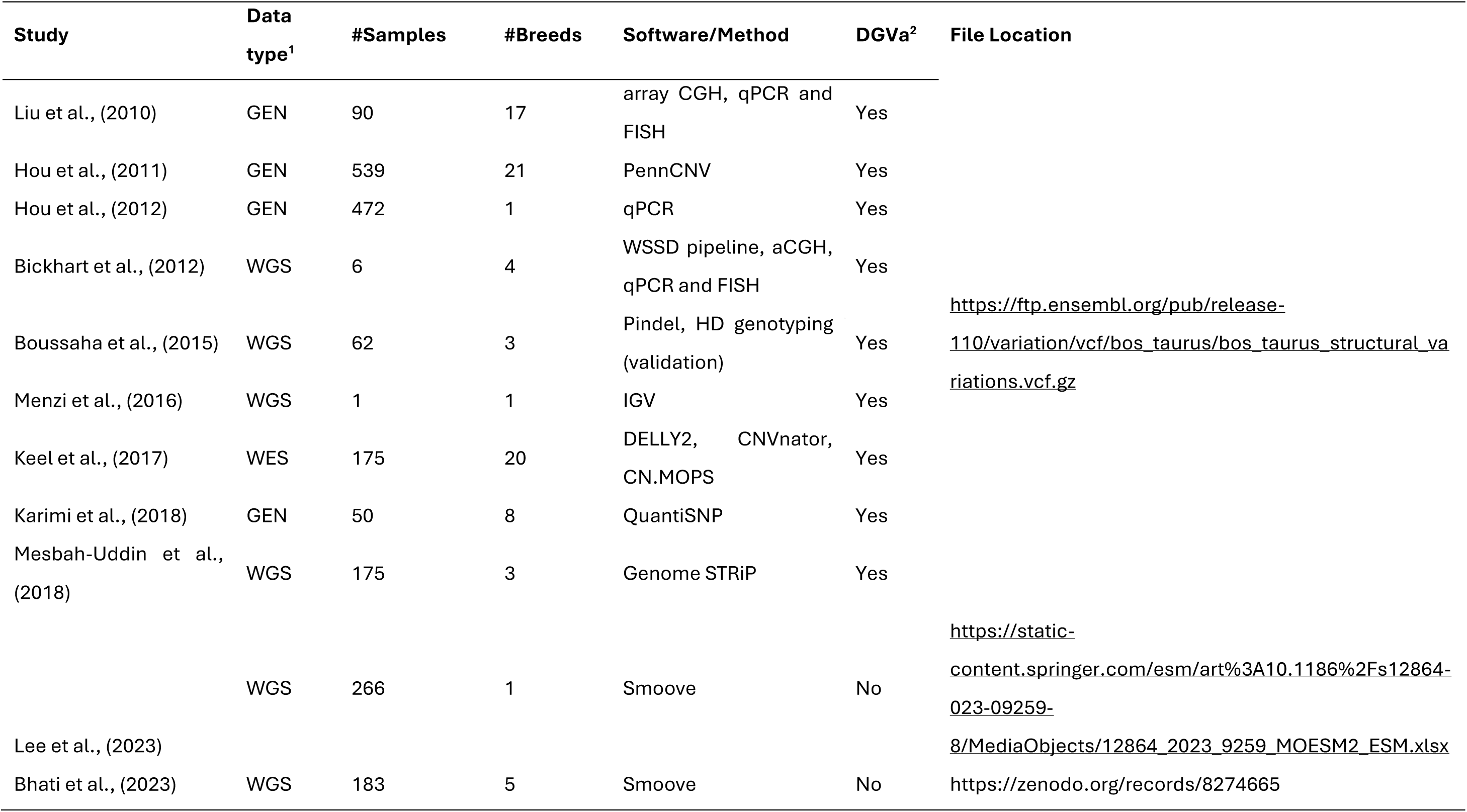
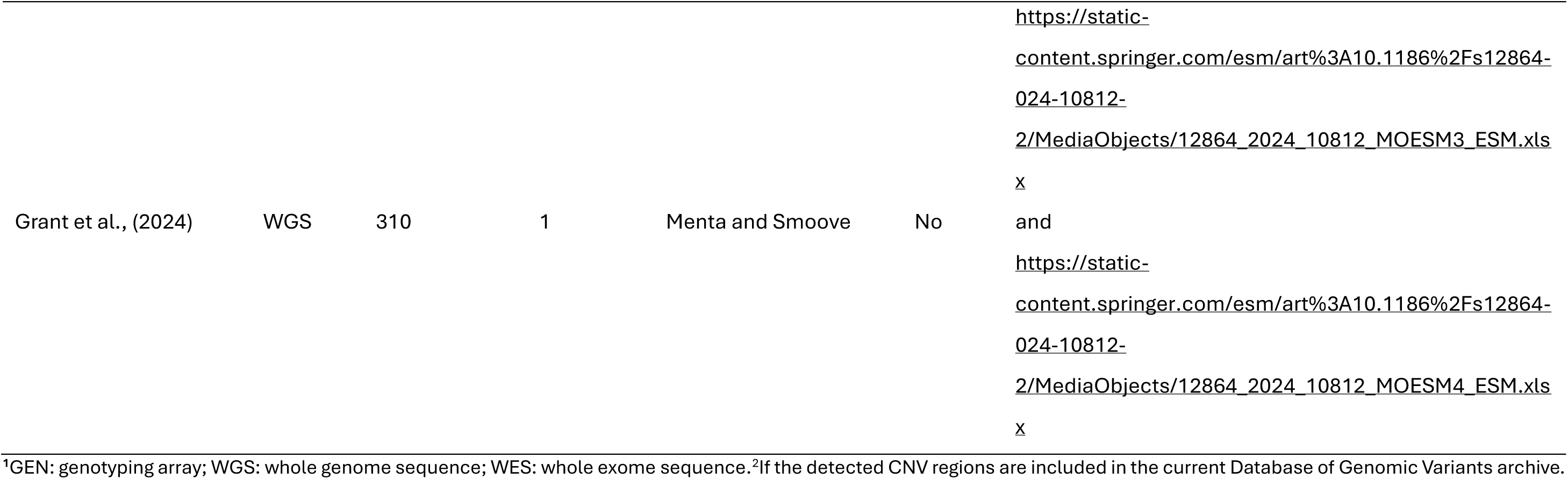
Summary of publicly available cattle CNV datasets used for comparison, including data type, sample size, breed composition, and analytical methods.

